# Elevated ozone concentration and nitrogen addition increase poplar rust severity by shifting the phyllosphere microbial community

**DOI:** 10.1101/2021.12.23.474070

**Authors:** Siqi Tao, Yunxia Zhang, Chengming Tian, Sébastien Duplessis, Naili Zhang

## Abstract

The tropospheric ozone and nitrogen deposition are two major environmental pollutants. Numerous studies have focused on the negative impacts of elevated O_3_ and the complementary effect of soil N addition to tree physiological characteristics. However, it was notoriously ignored of how elevated O_3_ with N addition affect tree immunity in face of pathogen infection, as well as of the important roles of phyllosphere microbiome community in host-pathogen-environment interplay. Here, we examined the effects of elevated O_3_ and soil N addition on poplar leaf rust (*Melampsora larici-populina*) severity of two susceptible hybrid poplars (clone ‘107’: *Populus euramericana* cv. ‘74/76’; clone ‘546’: *P. deltoides* ⊆ *P. cathayana*) in Free-Air-Controlled-Environment plots, besides, the link between *Mlp*-susceptibility and changes in microbial community was determined using Miseq amplicon sequencing. Rust severity of clone ‘107’ significantly increased under elevated O_3_ or N addition only, however, the negative impact of elevated O_3_ could be significantly alleviated when simultaneously conducting N addition, likewise, this trade-off was also found in its phyllosphere microbial *α*-diversity responding to elevated O_3_ and N addition. However, the rust severity of clone ‘546’ did not significantly differ in the cases of elevated O_3_ and N addition. *Mlp* infection altered microbial community composition and increased its sensitivity to elevated O_3_ assessed by significantly different abundance of taxa. Elevated O_3_ and N addition reduced the complexity of microbial community, which may explain the increased severity of poplar rust. These findings demonstrated that poplars need shifting phyllosphere microbial associations to optimize plant immunity in response to environmental changes.

**Importance:** Exploitation of the interaction mechanisms between host plants and pathogens is the essential basis in disease control. However, while much was known about the molecular determinants in pathogenesis process in the past decades, less is known about the role of nonpathogenic microbial community in plant-pathogen interaction, especially when some host plants are currently encountering severe environmental stresses, such as elevated ozone concentration and superfluous nitrogen addition. Thus, we targeted at the widespread and detriment rust disease (poplar-poplar rust) to dissect the influences of elevated ozone and nitrogen addition on rust disease severity and how phyllosphere microbial groups interacting with both poplars and rust pathogen under these biotic stresses. Our findings could be used to prescribe ecological information about poplar plantation in areas with high levels of ozone pollution and better understand the important roles of microbiome in plant heath.

## 1. Introduction

*Melampsora larici-populina* Kleb. (Basidiomycota, Pucciniales), the most devastating and widespread pathogen responsible for poplar foliar rust disease, caused severe treats to poplar plantations worldwide (1). In the wake of *Populus trichocarpa* genome sequencing (2) and the > 101 Mb genome sequencing of *M. larici-populina* (3), the molecular mechanisms underlying the binary interaction between poplar-poplar rust have been largely investigated in past decades (4–10). However, the current understanding of this disease must be revisited when encountering the complex nature of environmental changes, typically for variations in atmospheric composition which are expected to significantly aggravate the stress on plants (11).

Tropospheric ozone (O_3_) is considered as a major phytotoxic air pollutant, which enters plant tissues via stomata. Due to fast urbanization and industrialization, most forests in northern China were exposed to high concentration of O_3_ during the plant growing season (12). Acute doses of O_3_ exposure have series deleterious effects on plant growth and productivity, such as visible foliar injury, stomatal closure, reduction in photosynthesis, impairment of biomass and yield, and change in antioxidant capacity (13–14). While the tree growth responses have received much more attention, research concerning the elevated O_3_ on plant susceptibility to pathogens remains limited. Earlier studies indicate the acute increment of O_3_ could trigger defense against pathogens when used at appropriate concentrations. As soon as O_3_ penetrate plant tissues, it generates reactive oxygen species (ROS) such as superoxide anions and H_2_O_2_ by interacting with cellular components, leading to an alteration of several signal decades and other signal transduction pathway such as jasmonic acid (JA) and salicylic acid (SA) (15). The accumulation of SA and antioxidative defense by O_3_ exposure led to significantly decrease in the infectious intensity by obligate pathogens (16–18). However, the effects of O_3_ on plant susceptibility largely depend on the timing of exposure to O_3_. There is an evidence showing that the severity of wheat stem rust (*Puccinia graminis* f. sp. *tritici*) decreased by an exposure to O_3_ given 24 - 48h before inoculation but not by an exposure given after inoculation or before visible injury developed in host (19). As one of the largest plantations in China, poplar trees have to face the great challenge of long-period exposure to O_3_ pollution during the growing season. However, how poplars response to rust infection under the elevated O_3_ stress remains elusive.

In contrast to ozone, nitrogen (N) is the most essential inorganic nutrient that promotes plant growth. Adequate but not excessive amounts of nitrogen are required for efficient development of plants, such as regulation of metabolism, growth and resource allocation (20). However, anthropogenic activities (e.g., excessive uses of fossil fuels and fertilizers) have disequilibrated the N cycle in terrestrial ecosystems (21). Several studies have pointed out the negative effects of excessive N application on plant susceptibility to pathogens, especially increasing the severity of diseases caused by powdery mildew and stripe rust infection (22–23). These deleterious effects could attribute to the superfluous N content in leaf tissues, which provided a favorable environment for the pathogen growth and development (24–25). Generally, elevated O_3_ and N addition simultaneously affect plant growth in natural ecosystems (14). A recent study showed that N addition could decrease the accumulated O_3_ uptake by reducing stomatal conductance (26), thus it is crucial to clarify how this N addition - O_3_ flux trade-off influence the plant susceptibility

Phyllosphere, the aerial parts of plants, harbors hyperdiverse microbial communities with numbers ranging from 10^6^ to 10^7^ bacteria/cm^2^ (27). Microbiologists and plant pathologists have studied phyllosphere since mid-1950s (28), mostly because some foliar pathogens threaten plant health whereas others improve plant performance (29–30). Pioneer studies showed phyllosphere microorganisms played essential roles in hindering disease development through direct interactions (e.g., production of antibacterial or antifungal compounds) or indirect interactions (e.g., competition for foliar nutrients or alteration of plant physiology) with pathogens (29–30). Massive meta-sequencing during the last decade has fostered the study of phyllosphere microbial communities, providing a better understanding of non-culturable microorganisms (31). Systematic exploration of plant phyllosphere indicates the important roles in susceptibility to pathogens (32–35). However, the microbial community composition is variable as plants grew under various environments through recruiting different sets of microbes (36). A few studies revealed the decreased phylogenetic diversity of soil bacterial and archaeal communities of rice under elevated O_3_ concentration (37–39). However, the effects of elevated O_3_ on rhizosphere microbes could be very limited as the O_3_ concentration in soil is very low (40). Compared to below-ground microbial communities, phyllosphere microbiota colonized more extreme, stressful and changing environments as they interact directly with the dynamics of volatile organic compounds and atmospheric trace gasses (41). Even though the earlier report suggested O_3_-treated phyllosphere of rice harbors more variable bacterial communities (42), the structure and formation of fungal community under elevated O_3_ concentration are scarcely understood and the study on broadleaf forests remains largely under-explored. Most works examining the impact of N fertilization on microbiome composition and function focus on the rhizosphere (43–44), but not on plant leaves. Since the negative effect of elevated O_3_ on tree characteristics could be modified by N addition (26), information is needed on how this interaction impact on plant microbiome. Furthermore, the pivotal roles of phyllosphere microorganisms at the interface between O_3_ dynamics and N addition with plant disease has been largely neglected in the past. Hence, a more integrated recognition is needed that how phyllosphere microbe-microbe interactions influence plant immunity under elevated O_3_ and N addition.

The objectives of the present study are (1) to determine whether elevated O_3_ concentration, N addition and their interaction can modify poplar leaf rust severity; (2) to demonstrate how poplar phyllosphere microbial communities shift to defense rust infection under elevated O_3_ and N addition. We hypothesize that elevated O_3_ would cause a higher severity of *Melampsora larici-populina* (*Mlp*) -infection and N addition would alleviate the negative effect of elevated O_3_ on the ability of poplar defensing the *Mlp-*infection. We also expect that *Mlp*-infection would break down the stability of phyllosphere microbiome and their sensitivity responding to elevated O_3_ and N addition. To test the hypotheses, we selected two widespread hybrid poplars (clone ‘107’: *Populus euramericana* cv. ‘74/76’and clone ‘546’: *P. deltoides* ⊆ *P. cathayana*) which are planted in Free-Air-Controlled-Environment (FACE) plots and treated with elevated O_3_ and N addition. Results will provide pioneering insights into understanding how poplar respond to rust infection under elevated O_3_ and N addition and the potential roles of phyllosphere microbial community to play in this process.

## 2. Materials and Methods

### 2.1 Experiment site

The experiment was conducted at YanQing district, northwest of Beijing, China (40°47“N, 116°34E, elevation 485 m a.s.l). The region has a continental monsoon climate type. The annual mean temperature is 11.8 °C and the warmest month was July with a mean temperature of 24.5 °C. The average annual precipitation is 550 mm with about 44% falling between June and September (45).

### 2.2 Ozone fumigation treatment and nitrogen addition

An open-air O_3_ enrichment system in each Free-Air-Controlled-Environment (FACE) plot was used for O_3_ fumigation. The treatments were ambient ozone concentration (A-O_3_) and elevated ozone concentration (E-O_3_) (targeted at ambient O_3_ ⊆ 1.5) with four replicate plots of each treatment. Four E-O_3_ plots were separated from others by at least 70 m to avoid cross-contamination. The quantity and direction of the O_3_ release was controlled with an O_3_ monitor (Thermo Electron 49i, Thermo Scientific Co., USA) and data logger-controller (Campbell CR 10X, Campbell Scientific Co., USA), anemometer and wind vane. The O_3_ fumigation ran from May to October since 2018. For more details of the O_3_ fumigation system, see (46). Two hybrid poplars ‘107’ (*Populus euramericana* cv. ‘74/76’) and ‘the clone ‘546’’ (*P. deltoides* ⊆ *P. cathayana*) from the Chinese Academy of Forestry Sciences were used in this study for their differences in ozone sensitivity (14). The seedlings were grown under ambient air and then manually transplanted into the A-O_3_ and E-O_3_ plots. Each plot was split into two subplots in accordance with two poplar varieties. Half of the trees in northern subplots of A-O_3_ and E-O_3_ plots were supplied with ammonium nitrate solution every month to sum up to a total amount of N of 60 kg N ha^-1^ yr^-1^ (N_60_) while the remaining trees were treated without N addition (N_0_). As such, the control without elevated O_3_ treatment (A-O_3_), elevated O_3_ (E-O_3_), N addition (N_60_), without N addition (N_0_) and their combined treatments were involved in this experiment.

### 2.3 Evaluation of poplar foliar rust severity

During September 26^th^-27^th^ in 2020, when wild poplars were heavily infected by *M. larici-populina*, three *Mlp*-infected ‘107’ poplars and three *Mlp*-infected ‘546’ poplars were randomly selected from N_0_ and N_60_ subplots in four A-O_3_ and four E-O_3_ plots, respectively (Fig.S1). For each selected tree, more than ten *Mlp*-infected leaves with the total number of 1125 images were captured for rust severity quantification by analyzing the percentage of the abaxial leaf surface covered by uredinia (Table S1). A grid ruler was prepared for size calibration for all images. The leaf area and the number of uredinia were measured by the image analysis software Image-Pro Plus v 6.0 (Media Cybernetics, L. P., Silver Spring, MD) (Fig. S2). Firstly, we calibrated the geometry size of leaf image with ‘Spatial Calibration’. Then, we outlined along the edge of the leaf with the AOI (area of interest) tool. Select area as measurements and covert the outlined leaf profile to a measure object using ‘Convert AOI(s) To Object (s)’ menu. The area data of measured leaf could be accessible in ‘Measurement data’. Select the outlined leaf profile again, using the color separation method that based on the color histogram to select uredinia, choose ‘Measure objects’ and ‘Apply Filter Ranges’ respectively, set at ‘8-Connect’, ‘smoothing=25’, ‘fill holes’ and ‘convex hull’ and then click ‘Count’, the results of In range count are the number of uredinia in ‘select leaf’. Finally, artificially adjust the uredinia numbers compared to images to avoid mistaking. The severity of poplar foliar rust disease was evaluated by calculating the ratio of uredinia numbers and leaf area (uredinia/cm^2^).

### 2.4 Leaf samples collection, DNA extraction and Illumina amplicon sequencing

Three healthy (no uredinia on leaf) and three *Mlp*-infected leaf samples were synchronously collected from N_0_ and N_60_ treatments in each A-O_3_ plot and E-O_3_ plot, respectively for the ‘107’ clone and the ‘546’ clone. Each sample was then divided into two: one stored at 4 °C for chemical analysis and the other one stored at -80 °C for DNA extraction. Ten leaf discs of 1.2 cm diameter from one leaf were collected and dried until constant weight at 70 °C and then the dry mass of the discs was measured to calculate the leaf mass per area (LMA). Leaf samples were dried out at 70 °C for 96 h, finely grinded in mortars to estimate the total organic carbon (C) and total nitrogen (N) with CHNOS Elemental Analyzer (vario EL III, CHNOS Elemental Analyzer; Elementar Analysensysteme GmbH, Langenselbold, Germany).

Leaf genomic DNA was extracted using a DNA secure Plant Kit (TIANGEN, Beijing, China). The concentration of DNA extracts was determined using the NanoDrop 2000 UV-vis spectrophotometer (Thermo Scientific, Wilmington, USA), and the quality of DNA extracts were examined using 1% agarose gel electrophoresis. The primer pairs 799F (5’-AACMGGATTAGATACCCKG-3 ’)/1392R (5’-ACGGGCGGTGTGTRC-3 ’) and the primer pairs 799F (5’-AACMGGATTAGATACCCKG-3 ’)/ 1193R (5’-ACGTCATCCCCACCTTCC-3 ’) were used to amplify the V5-V7 region of the bacterial 16S rRNA gene by a nested PCR. The primer pairs ITS3F (5’-GCATCGATGAAGAACGCAGC-3’)/ITS4R (5’-TCCTCCGCTTATTGATATGC-3’) were used to amplify the ITS2 region of fungal ITS gene. The PCR amplification was performed as follows: initial denaturation at 95 °C for 3 min, followed by 27 cycles of denaturing at 95 °C for 30 s, annealing at 55 °C for 30 s and extension at 72 °C for 45 s, and single extension at 72 °C for 10 min, and end at 10 °C. The PCR mixtures contain 5 × TransStart FastPfu buffer 4 μL, 2.5 mM dNTPs 2 μL, forward primer (5 μM) 0.8 μL, reverse primer (5 μM) 0.8 μL, TransStart FastPfu DNA Polymerase 0.4 μL, template DNA 10 ng, and finally ddH2O up to 20 μL. PCR reactions were performed in triplicate. The PCR product was extracted from 2% agarose gel and purified using the AxyPrep DNA Gel Extraction Kit (Axygen Biosciences, Union City, CA, USA) according to manufacturer’s instructions and quantified using Quantus™ Fluorometer (Promega, USA). The qualified PCR products were mixed, and paired-end sequenced on an Illumina MiSeq PE300 platform (Illumina, San Diego, USA) according to the standard protocols by Majorbio Bio-Pharm Technology Co. Ltd. (Shanghai, China). The raw sequences are available in the NCBI Sequence Read Archive (SRA) database with the accession code PRJNA776974.

### 2.5 Bioinformatics and statistical analysis

All paired rRNA amplicon sequencing raw reads were processed via QIIME2 v2020-6 (47). The raw reads were imported into QIIME2 manually using the “qiime tools import” command. The quality trimming, denoising, merging and chimera detection were done using the plugin “qiime dada2 denoise-paired” in DADA2 (48) as implemented in QIIME2 v2020-6, the “-p-trim-left-f” and “-p-trim-left-r” parameters were set at 0 and the “-p-trunc-len-f” and “-p-trunc-len-r” parameters were set at 300 for bacteria and 300 for fungi, respectively, after reviewing the “Interactive Quality Plot tab” in the “demux.qzv” file. The α- and β- diversity analyses were conducted through the “core-metrics-phylogenetic” method in the q2-diversity plugin with the setting of “-p-sampling-depth” at 4529 for bacteria and 1851 for fungi, according to the “Interactive Sample Detail” in the “table.qzv” file. The bacterial AVSs were taxonomically classified using the qiime2 v2020-6 plugin “qiime feature-classifier classify-sklearn” with the pre-trained Naïve Bayes Greengenes classifier trimmed to the V5-V7 region of the 16S rDNA gene. The fungal ASVs were analyzed by UNITE classifiers against the UNITE reference database. Weighted Unifrac principal component analysis (PCoA) was used to assess the *β*- diversity across different treatments, followed by the significance test by permutational multivariate analysis of variance (PERMANOVA; 49). The Spearman’s correlation coefficients among Nitrogen addition, ozone concentration, leaf properties, rust severity, fungal α-diversity and bacterial α-diversity were analyzed and visualized by R package MatCorPlot (50). To determine the effects of elevated O_3_, N addition and *Mlp*-infection on phyllosphere associations in the two clones of poplar, the underlying co-occurrences between bacterial and fungal taxa were depicted through network analysis using the R library igraph (51). The network analysis was performed at the class level to reduce the complexity of calculation as well as to ensure the accuracy of taxonomic information. Data filtering was performed prior to network construction in that only highly abundant ASVs that were in the top 10% in terms of relative abundance across all samples were reserved to mitigate the random variances (52). The resulting correlations were then imported in Gephi software (53) and visualized by the Frucherman Reingold algorithms, the topology property parameters of the network the clustering coefficient, network density and modularization were calculated automatically in Gephi.

## 3. Results

### 3.1 Combined effects of elevated O_3_ concentration and N addition on foliar rust severity of two poplar clones

The elevation of O_3_ concentration and N addition had significant effects on rust severity for the ‘107’ clone, it while did not influence that of the ‘546’ clone (Fig.1). For the ‘107’ clone, the rust severity significantly increased (Wilcoxon test, *p* < 0.005) under elevated O_3_ concentration (E-O_3_ plots) compared to ambient O_3_ concentration (A-O_3_ plots). In four A-O_3_ plots, N addition also could significantly increase the rust severity, however, in four E-O_3_ plots, the combined effects of O_3_ concentration elevation and N addition led to significant decreases in rust severity compared to the single effect of elevated O_3_. For the ‘546’’ clone, there were no significant differences of rust severity observed between A-O_3_ and E-O_3_ plots. Besides, N addition did not alter the rust severity of the ‘546’’ clone in both A-O_3_ and E-O_3_ plots (Fig. 1).

**Fig. 1.**
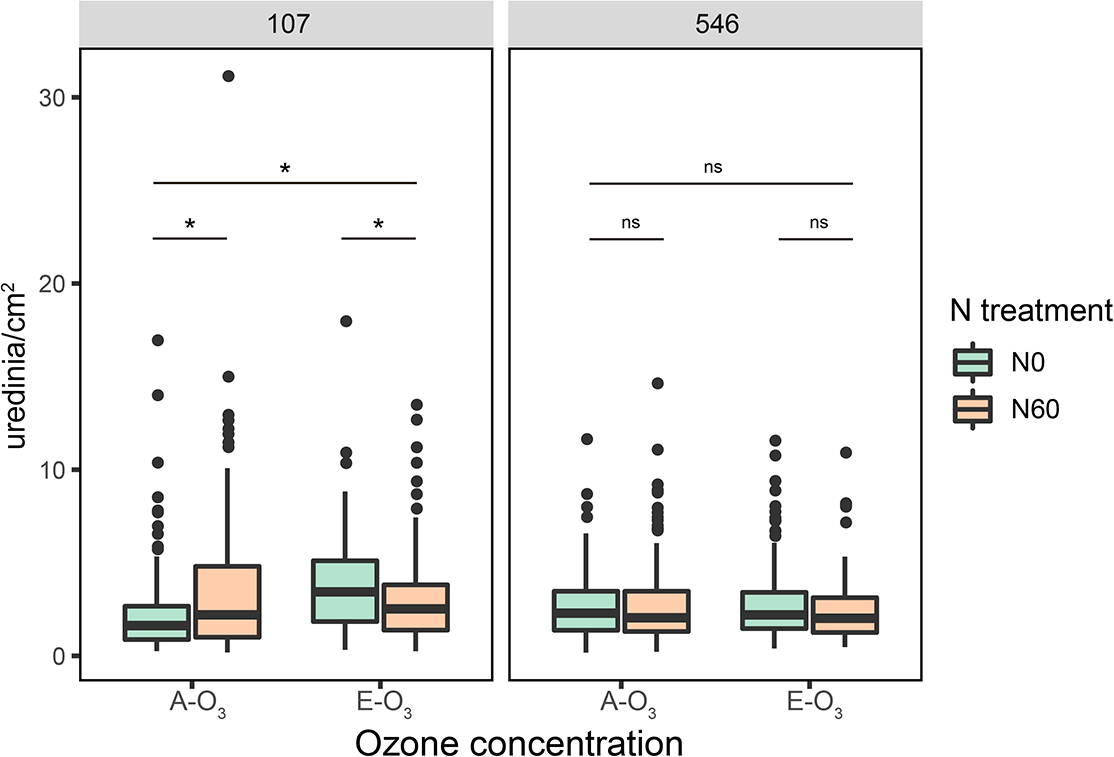
Rust severity of poplar foliar rust in four ambient ozone concentration FACE plots (A-O_3_) and four elevated ozone concentration FACE plots (E-O_3_) under two N treatments (N0 = no addition of nitrogen, N60 = addition of 60 kg/ha nitrogen every month). The significant differences between treatments at the 0.05 probability were indicated as the asterisk (*) according to the two-tailed Wilcoxon test; ns: not significant.

### 3.2 The relative abundance of phyllosphere fungal and bacterial species under elevated O_3_ concentration and N addition

Leaf samples used for estimating phyllosphere microbial composition characterization were collected from eight FACE plots (See “Methods” section). After the MiSeq PE300 high-throughput sequencing, a total of 7,570,222 and 8,148,246 raw reads from 16S rRNA and ITS were identified among 64 samples. After quality filtering, denoising, merging and chimera screening processes, 6,966 and 663 amplicon sequence variants (ASVs) were obtained for bacteria and fungi, respectively. Because fungal ITS primers also target host plant DNA, 155 ASVs (23.38% of total ASVs) were assigned to Viridiplantae, of which 94% were annotated as *Populus deltoides* (Table S2). After removing sequences affiliated to Viridiplantae, 508 remained ASVs were identified to fungi (Table S3). All ASVs derived from 16S rRNA were assigned to Bacteria (Table S3). Despite some amplicons were assigned to the plant, the sequencing depth was high enough to capture the majority of observed ASVs (Fig. S3). The dominant bacterial Phyla were Proteobacteria (68.1%), Actinobacteria (20.0%) and Thermi (9.6%). At the Class levels, bacterial ASVs classified the dominant class was Αproteobacteria (28.6%) followed by Actinobacteria (13.0%) and Gammaproteobacteria (9.3%), respectively (Fig. 2a). The three most dominant fungal groups at the genus level were *Phyllactinia* (8.60%), *Peyronellaea* (2.82%) and *Cladosporium* (1.68%), respectively (Fig. 2b).

**Fig. 2.**
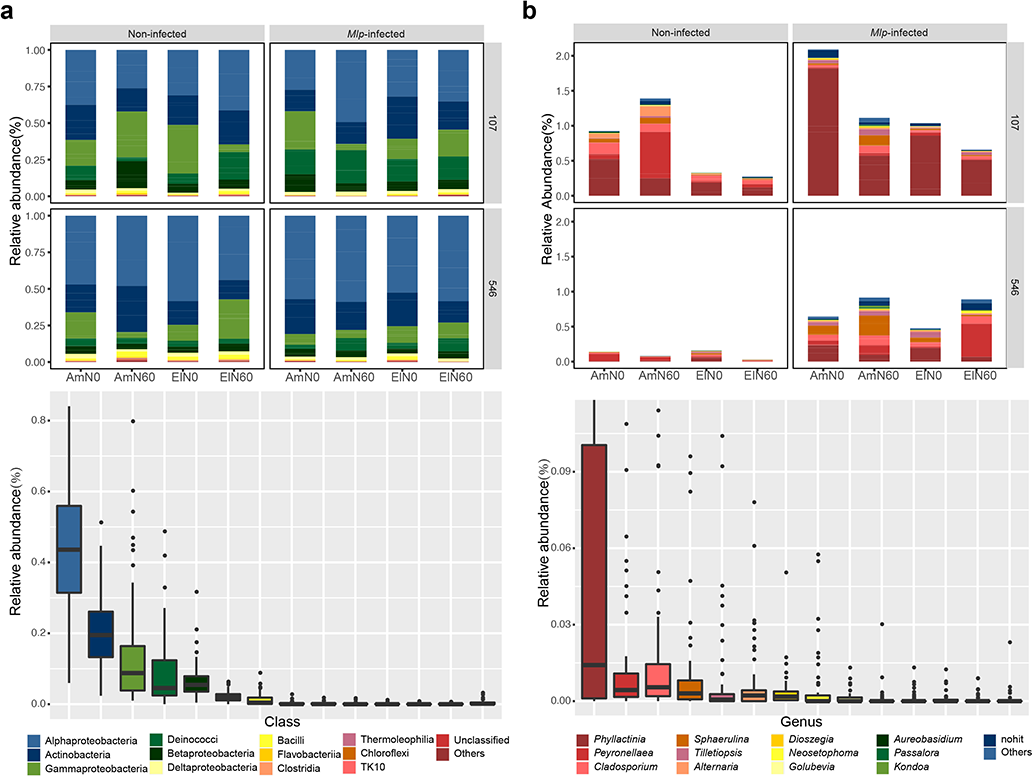
Taxonomic structures of phyllosphere bacterial microbiota at the class level (a) and fungal microbiota at the genus level (b). Only the 12 families with the largest mean relative abundance are shown.

Taxa significantly differed at the family level were identified using the generalized liner models (GLM, Table S4). For this analysis, Methylobacteriaceae and Aurantimonadaceae were significantly enriched in the ‘546’ clone, while Enterobacteriaceae, Microbacteriaceae, Deinococcaceae, Oxalobacteraceae and Pseudomonadaceae were significantly more abundant in the ‘107’ clone. We examined the differentially abundant bacterial taxa in *Mlp-*infected 107 leaves in A-O_3_ plots without N application and found three taxa significantly decreased (*p* < 0.01, FDR corrected), including Pseudonocardiaceae, Erythrobacteraceae and Bdellovibrionaceae. Four taxa significantly decreased (*p* < 0.01, FDR corrected) in *Mlp*-infected 546 leaves in A-O_3_ plot without N application: Corynebacteriaceae, Planococcaceae, Weeksellaceae and Propionibacteriaceae. In *Mlp*-infected leaves of the ‘107’ clone, elevated O_3_ significantly increased the bacterial groups within Corynebacteriaceae and Dietziaceae, while N addition did not significantly affect bacterial abundance. In *Mlp*-infected leaves of the ‘546’ clone, both elevated O_3_ and N addition had no impacts on bacterial abundance.

For fungi, we found 5 genera which were significantly differentially abundant (*p* < 0.01, FDR corrected) between poplar varieties. Of these, genera of *Phyllactinia*, *Cladosporium*, *Alternaria*, *Golubevia* were higher in the ‘107’ clone and genus of *Kondoa* was higher in the ‘546’ clone. For the clone of ‘107’, *Phyllactinia* and *Tilletiopsis* were significantly abundant in *Mlp*-infected leaves and *Peyronellaea*, *Cladosporium*, *Alternaria* and *Passalora* were significantly reduced in *Mlp*-infected leaves. For the ‘546’ clone, *Phyllactinia*, *Sphaerulina* and *Tilletiopsis* significantly increased in rust-infected leaves, while *Peyronellaea*, *Golubevia* and *Kondoa* significantly reduced. In *Mlp*-infected 107 leaves, *Alternaria* significantly increased under elevated O_3_. For *Mlp*-infected leaves of the ‘546’ clone, both elevated O_3_ and N addition had no significant impacts on fungal community structure (Table S5).

### 3.3 The variations of bacterial and fungal α-diversity under elevated O_3_ concentration and N addition

The differences in Shannon index, the representative of *α*- diversity, across different poplar clones, leaf condition, elevated O_3_ and N treatments were analyzed for both bacteria (Fig. 3a) and fungi (Fig. 3b). Using the Kruskal-Wallis rank sum test, we found no statistically significant differences in *α*- diversity across different conditions for bacterial community. As to phyllosphere fungi, their α- diversity in *Mlp*-infected leaves for the ‘546’ clone showed significantly higher (*p* < 0.05) than that of healthy leaves. On the contrary, fungal α- diversity of *Mlp*-infected and healthy leaves did not significantly differ in the clone ‘107’. Notably, the bacterial *α*- diversity in *Mlp*-infected leaves showed similar patterns in two poplar clones: increased with N addition in A-O_3_ but decreased with N addition in E-O_3_. However, the fungal *α*-diversity showed distinct patterns in *Mlp*-infected leaves of two poplar clones, which increased with elevated O_3_ and N addition in the ‘107’ clone but decreased under elevated O_3_ and N addition in the ‘546’ clone.

**Fig. 3.**
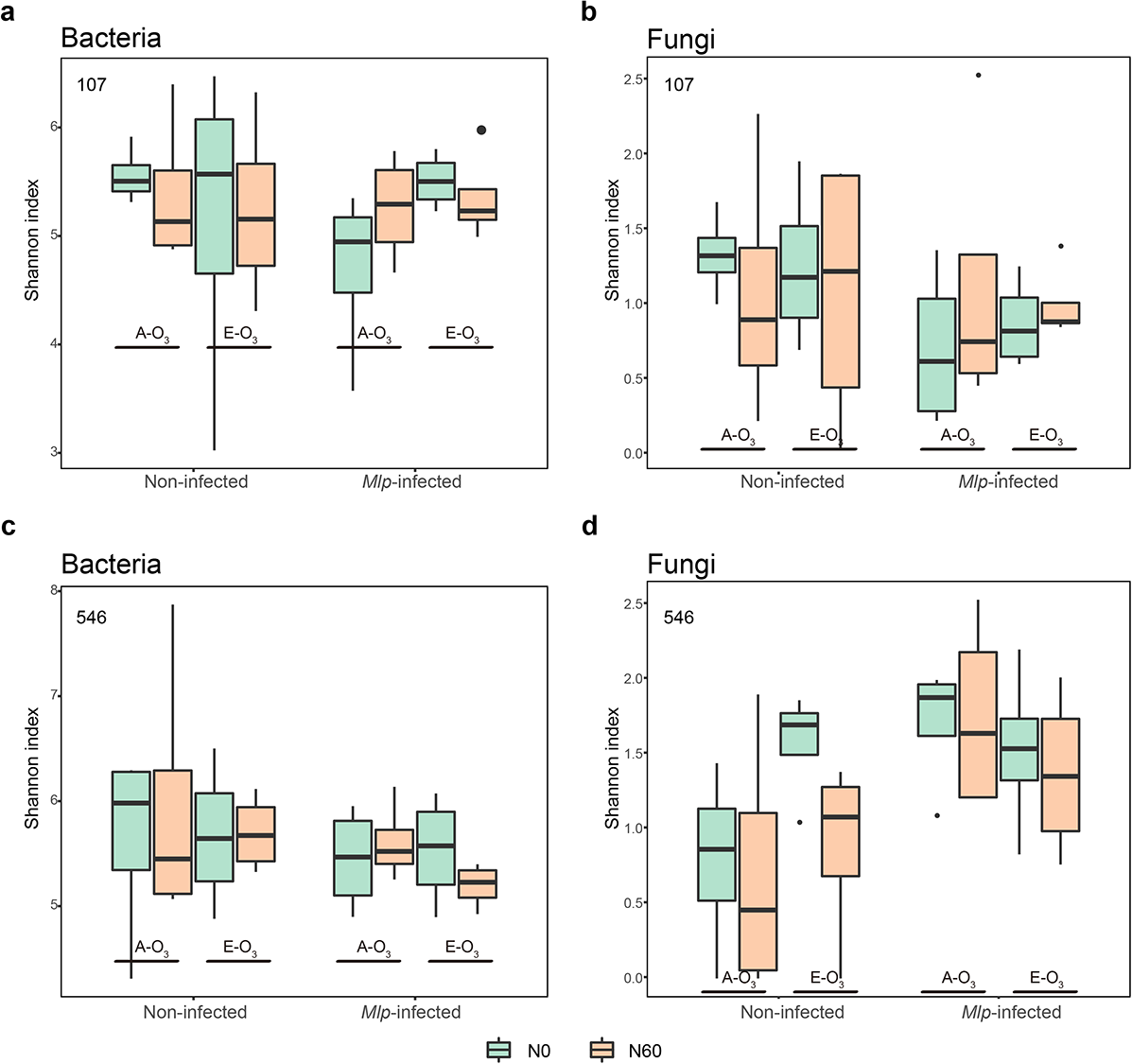
Shannon indices of phyllosphere communities of non-infected leaves and *Mlp*-infected leaves of the clone ‘107’ (a, b) and the clone ‘546’ clone (c, d) from ambient ozone concentration plots (A-O_3_) and elevated ozone concentration plots (E-O_3_) with nitrogen addition (N60) and without nitrogen addition (N0). Box plots showed the range of estimated values between 25% and 75%, the median, the minimum and the maximum observed values within each dataset.

### 3.4 The variation of bacterial and fungal *β*-diversity under elevated O_3_ concentration and N addition

The Weighted Unifrac distance matrices for both bacterial and fungal communities were calculated and visualized using PCoA analysis. As expected, the leaf condition (Non-infected vs. *Mlp*-infected) explained the largest part of variation in *β*-diversities of bacterial and fungal communities, 33.51% for bacteria and 79.15% for fungi, respectively (Fig. S4a, b). The poplar clone (‘107’ vs. ‘546’) was the second-largest indicator interpreting bacterial and fungal *β*-diversity variation (Fig. S4a, b). The clustering patterns of bacterial community under elevated O_3_ concentration and N addition were more pronounced for clone ‘107’ and which were clearly changed after infection of *Melampsora larici-populina* (Fig. 4a). Compared to bacterial community, fungal community was more affected by ozone elevation in both *Mlp*-infected leaves and non-infection leaves (Fig. 4b). PERMANOVA analysis also showed that poplar clone and leaf condition significantly influenced phyllosphere microbiome (*p* < 0.05). Like the PcoA result, poplar clone and *Mlp* infection more strongly influenced phyllosphere fungal community composition (F = 7.656, p = 0.002; F = 5.693, p = 0.008) than bacterial community composition (F = 6.661, p = 0.001; F = 2.702, p = 0.023) (Table 1). Notably, elevated O_3_ concentration significantly influenced the fungal community composition (F = 3.684, *p* = 0.037) rather than bacterial community composition (F = 0.288, *p* = 0.973), especially for the clone ‘107’ (F = 4.494, *p* = 0.020). Moreover, mental test was conducted for *Mlp*-infected leaves and the results suggested that nitrogen addition and leaf N content were the strongest environmental factors driving fungal and bacterial communities (Fig. 5) and rust severity was positively related to nitrogen addition and ozone elevation, which in addition, negatively related to fungal community alpha-diversity, bacterial Shannon diversity but positively relatively bacterial community richness and abundance (Fig. 5). Nitrogen addition and elevated O_3_ negatively related to both bacterial and fungal richness in *Mlp*-infected leaves (Fig. 5).

**Fig. 4.**
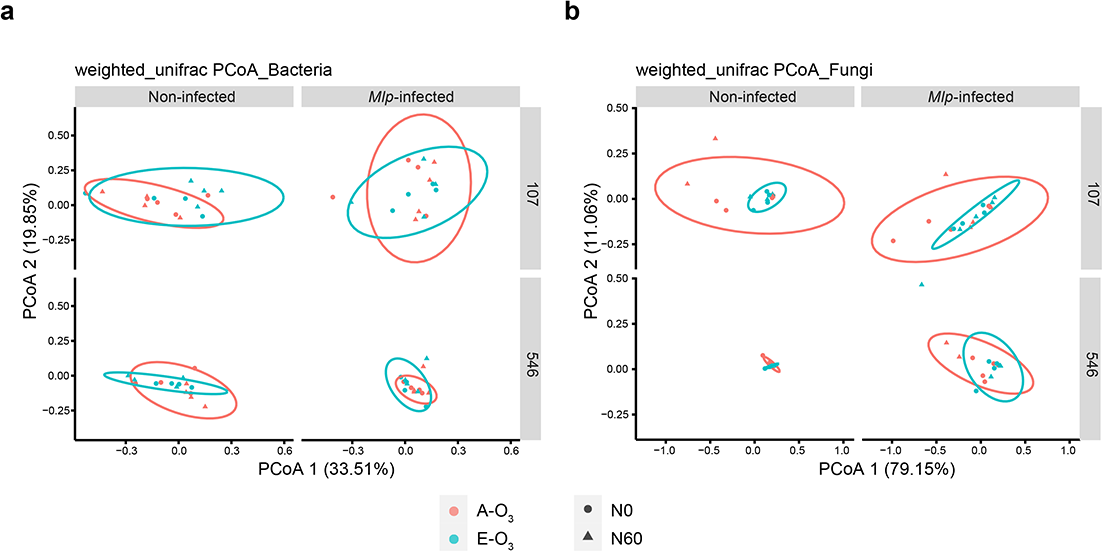
PCoA of bacterial and fungal communities using the weighted_unifrac distance for bacteria (a) and unweighted unifrac distance for fungi (b). samples are sorted for ozone concentration (A-O_3_ vs. E-O_3_) and nitrogen treatment (N0 vs. N60)

**Fig. 5.**
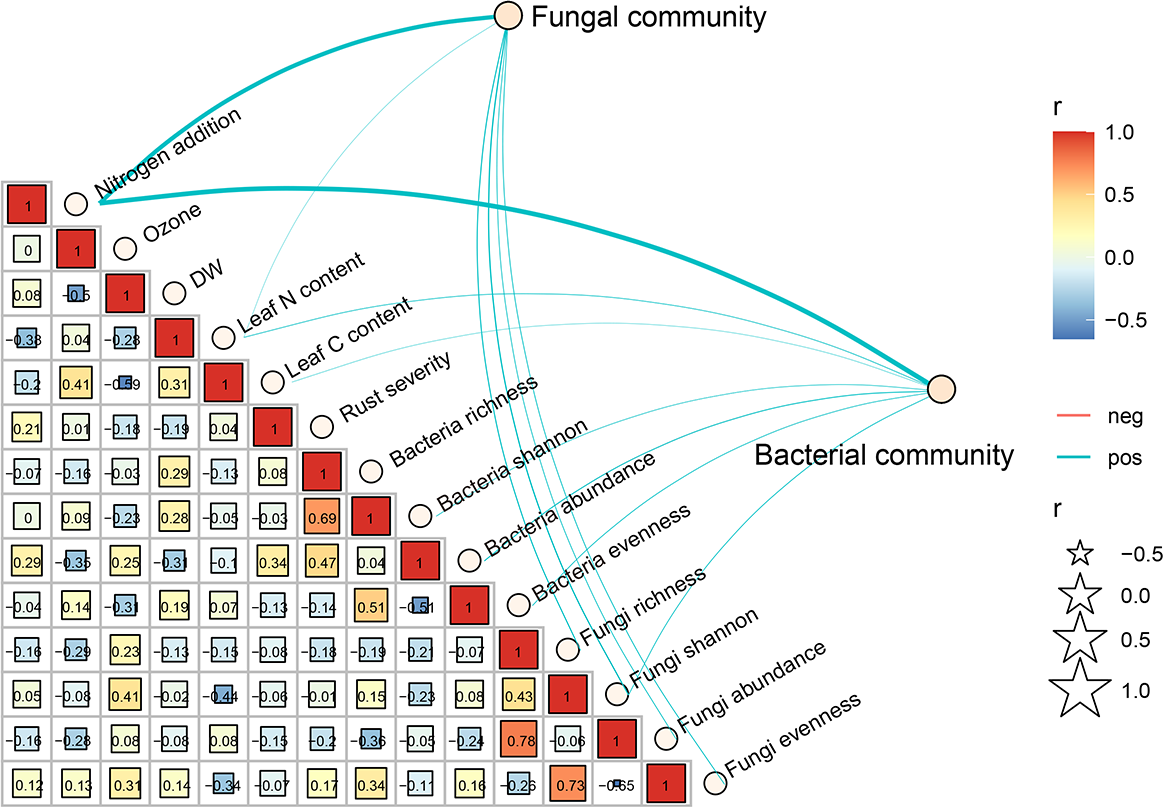
The mental test and Pearson correlation between α-diversity indices of *Melampsora larici-populina* infected poplar phyllosphere fungal and bacterial communities with nitrogen addition, ozone concentration (Ozone), leaf dry weight (DW), leaf nitrogen & carbon content and rust severity.

**Table 1.** The effects of polar clone, N addition, elevated O_3_ concentration and Melampsora larici-populina (Mlp) infection on the phyllosphere bacterial and fungal communities based on PERMANOVA analysis.

|  |  | Total |  |  |  | clone ‘107’ |  |  |  |  | clone ‘546’ |  |  |  |  |
| --- | --- | --- | --- | --- | --- | --- | --- | --- | --- | --- | --- | --- | --- | --- | --- |
|  |  | clone | N | O<sub>3</sub> | *Mlp* | N | O<sub>3</sub> | *Mlp* | N ×<br>O<sub>3</sub> | N ×<br>O<sub>3</sub> ×<br>*Mlp* | N | O<sub>3</sub> | *Mlp* | N ×<br>O<sub>3</sub> | N ×<br>O<sub>3</sub> ×<br>*Mlp* |
| Bacterial | *F* | 6.661 | 1.517 | 0.288 | 2.702 | 0.377 | 1.295 | 0.883 | 1.374 | 1.364 | 1.065 | 0.998 | 3.016 | 1.799 | 0.717 |
| community | *p* | **0.001** | 0.195 | 0.973 | **0.023** | 0.885 | 0.254 | 0.459 | 0.238 | 0.189 | 0.388 | 0.410 | **0.013** | 0.069 | 0.644 |
| Fungal | *F* | 7.656 | 0.776 | 3.684 | 5.693 | 1.081 | 4.494 | 2.756 | 1.410 | 0.534 | 0.846 | 0.337 | 6.223 | 1.013 | 0.435 |
| community | *p* | **0.002** | 0.422 | **0.037** | **0.008** | 0.312 | **0.020** | 0.085 | 0.231 | 0.602 | 0.406 | 0.762 | **0.003** | 0.347 | 0.587 |

### 3.5 Co-occurrence between poplar phyllosphere microbiomes

To disentangle the general effects of E-O_3_, N_60_ and *Mlp-*infection on phyllosphere microbiome co-occurrence patterns, we performed bacterial-fungal interkingdom network analyses of the clone ‘107’ and clone ‘546’ (Fig. 6a) and showed that the indices commonly used in assessing microbial network complexity (clustering coefficient, network density, number of nodes and number of edges) consistently decreased under N_60_, E-O_3_ and E-O_3_ + N_60_ treatments (Fig. 6b), indicating that poplar phyllosphere microbiome associations were less connected under these abiotic factors. With the addition of biotic stress (*Mlp*-infection), the microbial community of clone ‘107’ and ‘546’ exhibited totally different responses, the former presents a more complex association towards rust infection and the latter was just the opposite. Overall, the percentage of bacterial nodes in the networks of N_60_, E-O_3_, E-O_3_ + N_60_ and E-O_3_ + N_60_ + *Mlp* was reduced, and in contrast, the percentage of fungal nodes increased. Accordingly, the percentage of edges linking fungi-fungi and fungi-bacteria increased in the networks of N_60_, E-O_3_, E-O_3_ + N_60_ and E-O_3_ + N_60_ + *Mlp* compared to the control, whereas the percentage of edges between bacterial groups decreased in four treatments, these suggest a more active response in fungal community to (a)biotic stresses than bacterial community.

**Fig. 6.**
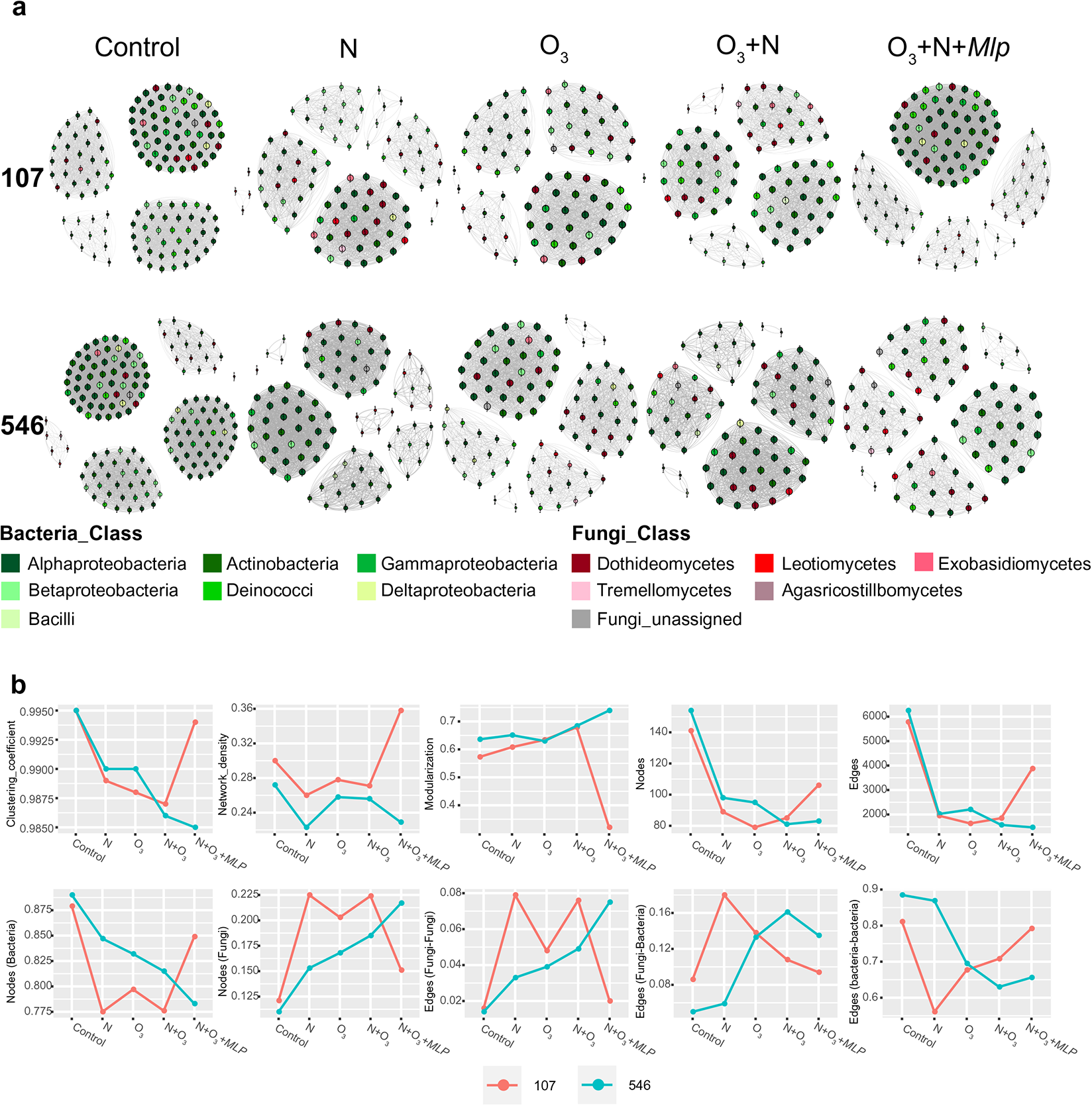
The co-occurrence networks (a) and trends of microbial network association indices (b) of phyllosphere microbiome for two hybrid poplars (‘107’ and ‘546’) in five conditions. No (a)biotic stresses (control), N addition (N), elevated O_3_ (O_3_), N addition with elevated O_3_ (N + O_3_), N addition, elevated O_3_ with *Melampsora-larici populina* infection (N + O_3_ + *MLP*)

## 4. Discussion

### 4.1 A trade-off of *Mlp*-susceptibility to elevated O_3_ and N addition for the ‘107’ poplar

In line with our hypothesis, we found exposure to elevated O_3_ throughout the growing season significantly predisposed the ‘107’ poplar to the infection of *M. larici-populina*, but not for ‘546’ poplar. Our results support the findings that growing-season-long exposures to enhanced O_3_ led to strongly positive effects on leaf rust of poplar (*Populus* sp.) caused by *Melampsora* species (18, 54). However, there were earlier studies in greenhouse chamber showing an acute dose of O_3_ could decrease the susceptibility of two eastern cottonwood (*Populus deltoides* Bartr.) clones to *Melampsora medusae* (17, 55). The opposite findings suggest the effect of elevated O_3_ on rust severity differs depending on the timing of exposure, even within rust pathogens (56). Short-time exposure to elevated O_3_ force the induction of ozone-related defense responses, such as increased transcription of genes from the phenylpropanoid pathway, PR genes and callose formation, and the priming of early senescence in leaves (57–58), which may explain the decreases in rust severity. After long-time exposures of O_3_, foliar surface topography, microroughness and physicochemical characteristics which determine the leaf surface properties and wettability were significantly changed (59), this O_3_-induced changes in leaf microenvironment may partially explain the significantly increases in rust severity of clone ‘107’.

Our results also verified the hypothesis that there is a trade-off of *Mlp*-susceptibility of poplar to elevated O_3_ and N addition, at least for ‘107’ clone. As many cases reported that inappropriate application of N fertilizer increases the severity of leaf rust diseases (24–25), our results showed N addition could exacerbate the ‘107’ poplar rust severity, possible mechanisms could be N addition-induced increase in leaf N content, providing nutrient resources for growth and reproduction of pathogenic fungi (60). However, N addition could alleviate the negative effect of elevated O_3_ on rust severity of clone ‘107’. According to a recent study showing N addition could limit the stomatal O_3_ uptake (26), we believe that this N addition - O_3_ flux trade-off could also influence the *Mlp*- susceptibility of poplar.

Clone ‘546’ is much more sensitive to O_3_ than clone ‘107’ (61). However, our results clearly showed that both elevated O_3_ and N addition did not significantly alter the *Mlp*-susceptibility of clone ‘546’. A recent study demonstrated that elevated O_3_ could reduce area-based leaf N concentration (N_area_) in poplars, which is positively related to photosynthetic parameters, and more O_3_ sensitive clone ‘546’ showed much greater reduction than clone ‘107’ (61). Therefore, we inferred that long O_3_ exposure have a negative impact on N allocation, furthermore, providing less nutrient for obligate biotrophs (*Mlp*). The interactive effects of O_3_ elevation and consequent N reduction may account for the insignificant change in rust severity of clone ‘546’.

### 4.2 The composition of poplar phyllosphere microbiome shifts under elevated O_3_, N addition and *Melampsora larici-populina* infection

Advances in culture-independent methods and next-generation sequencing technologies promote a better knowledge of composition of plant microbiome (62). By 16S rRNA and ITS sequences targeting, the structure features of the phyllosphere microbial communities of *Populus* clone ‘107’ and clone ‘546’ collected from sites with elevated O_3_ and N addition were characterized in this study. The dominant bacterial groups in the phyllosphere were Αlphaproteobacteria, Actinobacteria and Gammaproteobacteria, confirming that the most common groups of bacteria present in the phyllosphere are Proteobacteria, Bacteroidetes, Actinobacteria and Firmicutes (63). In our study, the composition of bacterial and fungal communities colonized in the phyllosphere of two poplar clones is different, with specific groups enriched. The cultivar-specific microbiome composition could possibly link to their phenotype and immunity (64–65).

It is well documented that the community of non-pathogenic microbes living in or on the leaves also influence the plant disease severity (33–34, 66) and diseased plants harbor altered microbiome compared with healthy plants (67). Our results indicated that *Phyllactinia* species dominated in two hybrid poplar phyllosphere and significantly increased after *M. larici-populina* infection. All ASVs assigned into *Phyllactinia* were further identified as *Phyllactinia populi*, which is a common foliar pathogen of *Populus* species in Asia (68), hence it is somewhat inevitable to see the poplar leaves developed with rust and powdery mildew in the field. A growing body of evidence supports essential roles of foliar fungi in disease modification (69–70). An inoculation experiment demonstrates that *Alternaria* and *Cladosporium* can reduce the severity of poplar rust disease as candidate pathogen antagonists (32). Furthermore, it has been reported that one *Alternaria* species hyperparasite in the urediniospores of *Puccinia striiformis* f. sp. *tritici* (71). Our molecular field surveys showed that *Peyronellaea*, *Alternaria* and *Cladosporium* presented a noticeable decrease in abundance in *Mlp*-infected leaves of clone ‘107’ or clone ‘546’. We postulate such changes in community composition may be due to the inability to compete with rust fungi when they successfully colonized in host plants (e.g., the competition of nutrients) or changes in phyllosphere microenvironments (e.g., chemical compounds and topography) hinder the development of these candidate antagonists.

Under ozone stress, plant emit specific volatile organic compounds to scavenge incoming ozone (72). The changes in carbon availability also affected the community composition of phyllosphere microorganisms (41, 73). However, elevated O_3_ only have significant effects on OTUs rather than higher taxonomy levels in the phyllosphere of rice (42), similar to the findings in our study that elevated O_3_ and N addition have little effect on the bacterial and fungal community composition at higher levels of taxonomic classification (family and genus). However, *Mlp*-infection raise the sensitivity of specific microbial groups in phyllosphere of clone ‘107’ which were significantly shifted responding to elevated O_3_, even at high taxonomic levels.

### 4.3 Elevated O_3_ and N addition lead to distinct responses in the phyllosphere microbial community diversity

The above-ground responses to elevated O_3_ and N addition in aspects of plant growth and photosynthesis have received much attention (74–75), the interactive effects of O_3_ and N on phyllosphere communities has been barely investigated. The host plant could directly affect the plant-associate microbial community by modifying the chemical features in surrounding environment (76). For instance, a recent study showed that N addition significantly decreased the rhizosphere soil bacterial α-diversity of the poplar clone ‘107’ and this negative effect could be alleviated by elevated O_3_ (77). Compared with rhizosphere, it is rarely known about how the role of abiotic factors in aerial tree surface, which is characterized as extremely poor in nitrogen and carbon sources and prone to rapid fluctuation (66, 78). Our studies characterized how N addition affect the phyllosphere microbiome community and found N could also negatively influence phyllosphere bacterial α-diversity of two hybrid poplar clones (‘107’ and ‘546’) and the gap narrows when O_3_ concentration elevated. For fungal community, N also decreased α-diversity for two clones, but the interactive effects of elevated O_3_ and N differ in two clones. The community from O_3_-treated rice phyllosphere was proved more diverse than those from control plants (42). However, our results presented totally different results that elevated O_3_ decreased α-diversity of phyllosphere bacteria and fungi of both clone ‘107’ and clone ‘546’. It was proposed that how elevated O_3_ impact the phyllosphere microbial diversity differs between monocotyledons and dicotyledon plants.

Our results verified that *Mlp*-infection could break down the balance of microbial community and influence their responses to elevated O_3_ and N addition. The microbial α- diversity of poplar phyllosphere microbiome decreased after the infection of *M. larici-populina*, likely due to high diversity supporting more mutualistic microbial interaction with plant immune systems to avoid pathobionts arising (65). Also, we surprisingly found that the bacterial and fungal communities in *Mlp*-infected phyllosphere of clone ‘107’ exhibited concurrent patterns in response to elevated O_3_ and N addition: increasing with elevated O_3_ and N addition whereas decreasing under their combined effects, in accordance with the patterns of rust severity. We suggest that the raises in α-diversity may be linked to specific microorganisms enriched responding to O_3_ elevation and N addition, which may affect the plant susceptibility through direct microbe-microbe interactions of and indirect interactions via nutrient competition, water-use efficiency and phytohormone production (79–80). But oppositely, the phyllosphere bacterial and fungal communities of the poplar clone ‘546’ exhibited completely opposite patterns under elevated O_3_ and N addition. We suppose that the effects of fungal and bacterial communities on *Mlp*-susceptibility neutralize with each other, potentially accounting for the insignificant changes in its rust severity.

The innate genetic traits of the plant (genotype and phenotype) could mediate leaf tissue chemistry and the lateral surface topology (e.g., roughness) that influences microbial immigration and emigration (65). Our PERMANOVA analysis performed on weighted_unifrac distances showed that the poplar clone was the major driver of variation in both bacterial and fungal communities, which may be due to the significant differences in morphological parameters between clone ‘546’ and clone ‘107’ (26). Besides, rust diseased-induced changes in the phyllosphere microbiome were also identified in this study, which consequently affect the plant-microbe and microbe-microbe interactions in the phyllosphere. The role of plant exudates in the reconstruction of phyllosphere communities and recruitment of beneficial microorganisms to resist the invasion of pathogens is of great concern in recent studies (81–82). Previous studies showed that infection by *Melampsora* species could induce flavonoid pathway-related genes (83–84) whereas identification of these plant metabolites in shaping the phyllosphere microbiome and immune-related regulatory mechanism remains great challenges. By comparing the effects of elevated O_3_ and N addition on rhizosphere soil microbiome community of poplar clone ‘107’, a study has shown that N addition may have a more direct effect on the belowground system than elevated O_3_ (77). Our finding suggests that elevated O_3_ may have a more direct effect on the aboveground system, as elevated O_3_ exert a significant impact on the composition of phyllosphere fungal community than N addition.

### 4.4 Effects of elevated O3, N addition and the combination of rust infection on co-occurrence of phyllosphere community

The plant host and its associated microorganisms interact dynamically to form a stable holobiont where the partners cooperate to increase the fitness (85). Hence, the functional capacity of a microbial community is not equal to the sum of its individual components, because microbial species interact with each other and form a complex network that has important implications for ecological processes and host adaptation (86). Distinct responses in microbial co-occurrence patterns were observed in response to elevated O_3_, N addition and the combination with *Mlp* infection as the number of edges, number of nodes and clustering coefficient decreased compared to the control. We interpret the decreased network size and complexity as a reduced community organization with weak interaction among the phyllosphere microorganisms. In contrast to the increased interaction network stability of rhizosphere microbial community under biotic and abiotic stresses in many cases (87–88), attenuate cooperation among phyllosphere microorganisms under elevated O_3_ and N addition is possibly driven by the inactive or dormant state of specific ozone-associated and nitrogen-fixing bacteria (41, 89). Indeed, although two hybrid poplars exhibited lower network complexity in the bacterial community, the percentage of fungal nodes and edges linking fungi to fungi showed high levels compared to the control. It is assumed that when threatened by pathogens, the multi-trophic interactions between kingdoms are disrupted, and the native microbial community must be restructured (90–91). We observed enhanced phyllosphere community organization in *Mlp*-infected leaves compared to non-infected leaves under elevated O_3_ and N addition. Under the combined effects of abiotic (elevated O_3_ and N addition) and biotic (rust fungi) stresses, plant commensal microbes that survive competition with diverse plant-associated microbes are more tightly connected than under single stress. Co-occurrence network analysis was also used to identify hub-microorganisms which are substantially more connected based on centrality measurements (92). However, identification of hub microorganisms which could exert strong direct and indirect effects on microbiome assembly and their functional roles in mediating between the plant-pathogen interactions under elevated O_3_ and N addition were underestimated in this study.

## 5. Conclusion

Overall, this study highlights the rust severity of one hybrid poplar (clone ‘107’) significantly increased under elevated O_3_ or N addition in Free-Air-Controlled-Environment (FACE) plots, while their interaction could diminish this negative effect. The phyllosphere microbiome of two poplar hybrids was dominated by specific microbial groups, and several taxa were significantly changed after rust infection. The bacterial α- diversity decreased with elevated O_3_ and N addition, irrespective of rust infection. For the clone ‘107’, the bacterial and fungal diversity in *Mlp*-infected phyllosphere showed different degrees of correlation with its rust severity. However, the trends of diversity for bacteria and fungi showed totally different pattern in the clone ‘546’, which may explain the insignificant changes in its rust severity. The study in phyllosphere microbial community composition observed across different conditions opens the possibility that host-specific traits are major drivers of variations followed by the biotic stress (*Mlp*-infection). Elevated O_3_ only had limited impacts on fungal community composition, while N addition had few influences on phyllosphere microbiome community. Finally, co-occurrence network analysis of phyllosphere microbiome indicates simplified microbial community under elevated O_3_, N addition and their combinations with *Mlp* infection, but the hub microorganisms that crucially connected with biotic and abiotic stresses as well as other microbes in network remains unknown, the following metagenome level analysis could be more informative and provide a functional view of those microbes.

## Acknowledgments

This research was supported by the Fundamental Research Funds for the Central Universities 2021ZY02 and China Postdoctoral Science Foundation under Grant No. 2021M690418.

## Author contributions

Siqi Tao: Investigation, Visualization, Writing-Original Draft, Funding Acquisition; Yunxia Zhang: Validation, Software; Chengming Tian: Conceptualization, Supervision; Sébastien Duplessis: Writing-Review & Editing; Naili Zhang: Conceptualization, Supervision, Writing-Review & Editing.

## Supplementary files

**Fig.S1.** The diagram of ambient ozone concentration (A-O_3_) plots and elevated ozone concentration (E-O_3_) plots. Each plot was divided into four subplots in accordance with two hybrid poplars (107: *Populus euramericana* cv. ‘74/76’; 546: *P. deltoides* ⊆ *P. cathayana*) and two treatments (N0 = no addition of nitrogen, N60 = addition of 60 kg/ha nitrogen every month). In each subplot, three rust-infected poplars and three non-infected poplars were randomly selected for subsequent analysis.

**Fig.S2.** The reduced illustration for quantifying the poplar leaf rust severity using Image-Pro Plus v 6.0.

**Fig.S3.** Rarefaction curves of sequence variants for each sample.

**Fig.S4.** PCoA of bacterial and fungal communities using weighted unifrac distance for bacteria (a) and unweighted unifrac distance for fungi (b).

**Table S1.** The dataset of poplar rust severity indices of 1125 Melampsora larici-populina infected poplar leaves collected from four ambient ozone concentration FACE plots (A-O_3_) and four elevated ozone concentration FACE plots (E-O_3_) with two N treatments (N0 = no addition of nitrogen, N60 = addition of 60 kg/ha nitrogen every month); the leaf_area (cm^2^) and numbers of uredinia were calculated using Image-Pro Plus v6.0; the dinia_per_cm^2^ representing the severity was calculated using the formula: numbers of uredinia / leaf_area (cm^2^).

**Table S2.** The taxonomic classification of bacterial and fungal amplicon sequence variants (ASVs).

**Table S3.** The count numbers of each amplicon sequence variant (ASV) across all samples.

**Table S4.** The generalized liner model (GLM) analysis results of bacterial taxa at the family level.

**Table S5.** The generalized liner model (GLM) analysis results of fungal taxa at the family level.

